# Group personality, rather than acoustic noise, causes variation in group decision-making in guppy shoals

**DOI:** 10.1101/2025.02.05.636607

**Authors:** Molly A. Clark, Ella Waples, Andrew N. Radford, Stephen D. Simpson, Christos C. Ioannou

## Abstract

Group living has essential fitness benefits for many species. While numerous studies have explored how environmental conditions impact collective movement, their impact on decisions made in a social context—a central component of group-living—is poorly documented. In this study, we assess how acoustic noise impacts group decision-making, cohesion and activity in fish shoals, using Trinidadian guppies (*Poecilia reticulata*) as a model species. Movements within a radially symmetric five-armed maze were measured using high-resolution trajectory data from video-tracking software. The behaviour of groups with and without continuous acoustic white noise were measured over a four-day testing period in a repeated-measures design. We found no significant change in swimming speed or group cohesion with additional acoustic noise. However, there was evidence for fewer following events (moves into already occupied arms) in the noise treatment compared to the control, but this additional noise had no effect on leadership attempts (moves into empty arms). We found strong evidence for consistent, repeatable differences between groups in all behavioural parameters indicating strong personality variation at the group level. Rather than environmental factors, these results provide evidence that consistent group-level differences dominate variation in collective behaviour, including group decision-making, in fish shoals.

## Introduction

Group living in animals can improve foraging, efficiency of movement and the avoidance of predators, with important fitness benefits (Krause & Ruxton, 2002; Ioannou, 2021). Many of these benefits are obtained through cohesive movement mediated by social interactions and self-organisation that allow for information transfer between individuals (Ioannou and Laskowski, 2023a). Group-living also has costs (Krause and Ruxton, 2002), and whether or not conformity is favoured by group members to maintain cohesion can depend on individual fitness requirements (Ioannou and Laskowski, 2023b).

Some of the benefits of group-living are provided by collective decision-making, which can enhance abilities to exploit resources and evade predation (Ioannou and Laskowski, 2023a). Group decisions can involve choices such as the direction of travel, where to forage, or where to nest (Conradt and Roper, 2005; Sumpter, 2006). Group decisions may be egalitarian, where decisions are equally shared among individuals, or involve leadership by a few individuals within a group, or even by a single individual (Conradt and Roper, 2005). Leadership is dependent on the ‘willingness’ of other individuals to follow, thus leadership attempts may be successful or unsuccessful depending on the actions of others in the group (King, 2010).

Fish in shoals are thought to use cues based on the location and movement of other individuals, rather than active signals, to mediate collective movement (Ioannou *et al*., 2011; Lemasson *et al*., 2018). Leaders emerge through positional changes within a group, with those at the front of a shoal having more influence over the direction of movement (Bumann and Krause, 1993; Krause *et al*., 2000). While studies of the mechanisms of collective decision-making are extensive, the impact of ecological context is less well known. Evidence that group decision-making can change with ecological conditions has been demonstrated in Trinidadian guppy (*Poecilia reticulata*) shoals, where Ioannou *et al*. (2017) found that group decision-making differed depending on whether individuals were caught from a low- or high-predation river. Guppies from high predation populations showed stronger differentiation into leader and follower roles, marked by a strong negative correlation between the number of leadership attempts and following events, compared to guppies from environments with lower predation pressure (Ioannou *et al*., 2017). The majority of studies testing how environmental stressors affect collective behaviour in fish shoals focus on collective motion, rather than group decision-making (Fisher *et al*., 2021). Therefore, there have been relatively few empirical studies of how abiotic conditions affect the dynamics of group decision-making. Chamberlain and Ioannou (2019) found that with experimentally induced water turbidity, three-spined sticklebacks (*Gasterosteus aculeatus*) became more independent in their decision making, shifting their behaviour away from that of the group, due to the visual constraints imposed by turbidity.

Anthropogenic noise pollution is a growing concern, particularly in aquatic environments (Slabbekoorn *et al*., 2010; Duarte *et al*., 2021). Human activities including ship traffic, construction, tourism and recreation have increased significantly over the past century, and with them so has anthropogenic sound (Peng *et al*., 2015). Due to the relatively high density of water compared with air, noise travels further and faster underwater, meaning the impacts of noise pollution can be more widespread (Slabbekoorn *et al*., 2010; Duarte *et al*., 2021). Animals use acoustic signals for communication and gather additional acoustic information about their environment. Anthropogenic noise can thus mask auditory reception of important stimuli, decreasing the signal-to-noise ratio (Slabbekoorn *et al*., 2010). Additionally, noise can distract individuals from important tasks, such as foraging (Purser and Radford, 2011; Voellmy *et al*., 2014a), and cause direct physiological stress (e.g. increasing cortisol levels: Smith *et al*., 2004; Wysocki *et al*., 2006).

Impacts of acoustic noise pollution can have knock-on effects on social behaviour (Fisher *et al*., 2021). Cetaceans have been found to alter their calls, in frequency (Parks *et al*., 2007) and duration (Foote *et al*., 2004), as a result of boat noise, demonstrating that acoustic noise can cause marine mammals to modify their acoustic communication. Noise-induced distraction and stress can also disrupt collective behaviour by impairing the effective use of information through cross-modal effects (Morris-Drake *et al*., 2016). Noise has been shown to impact collective movement negatively by reducing the coordination and cohesion of shoals (Sarà *et al*., 2007; Herbert-Read *et al*., 2017). However, whether noise disrupts the decision-making processes of groups has yet to be addressed.

We aimed to investigate how acoustic noise impacts individual behaviour, group cohesion and collective decision-making in shoals of Trinidadian guppies. Similarly to Ioannou *et al*. (2017), we recorded groups of guppies swimming in a five-arm radially symmetrical maze whilst experiencing an acoustic playback of either a recording from their housing tank (i.e. a control treatment) or this same recording with white noise overlaid to simulate acoustic noise pollution. By using high-resolution tracking data obtained by overhead video footage, we explored how the individual (i.e. speed and exploration) and group behaviours (cohesion and collective decision-making) of fish shoals changed with noise disturbance. To assess the consistency of behavioural responses we used a repeated-measures design, where the same groups of fish were tested across multiple days in both treatments. This approach allowed us to test whether changes in behaviour due to noise disturbance were consistent over time, and to explore group-level personality traits. Consistent behavioural responses across repeated trials would be evidence of group-level personality (Planas-Sitjà *et al*., 2015; Salazar *et al*., 2015; Jolles *et al*., 2018; MacGregor and Ioannou, 2021).

We hypothesised that additional acoustic noise would reduce overall group cohesion and exploration behaviour. Previous studies have shown that over repeated trials, animal groups tend to acclimate to experimental arenas, leading to reduced cohesion and altered exploratory behaviour (Miller and Gerlai, 2012; MacGregor and Ioannou, 2021; Allibhai *et al*., 2023). Therefore, we also expected group cohesion to reduce over the days of testing, while individual speed and group exploration may increase, as the environment becomes more familiar. In terms of collective decision-making, we expected that added noise would affect leadership dynamics, with a smaller proportion of leadership attempts being successful due to fewer following events. This could result from distraction or stress making potential followers less responsive to social cues.

## Materials and Methods

### 1 Ethics statement

The experiment was carried out at the University of Bristol aquarium and was approved by the university’s Animal Welfare and Ethical Review Body (UIN/17/060 and UIN/17/075). Methods followed ASAB Guidelines for the treatment of animals in behavioural research. Individuals were exposed only to two 20-minute additional-noise treatments to minimise stress. After the experiment, fish were monitored to ensure they were healthy, and housed in preparation for future experiments.

### 2 Study species

Guppies were originally collected from a high-predation site on the Guanapo river in Trinidad, West Indies, in April 2019. They were exported to the John Krebs Field Station, University of Oxford, and reared for a minimum of three generations with a controlled breeding plan to prevent inbreeding and ensure genetic diversity was maintained. Individuals from the third generation were transported by car for approximately 1.5 hours in sealed plastic bags, with 1 third water and 2 thirds oxygen, within insulated containers and then housed at the University of Bristol aquarium in December 2020. All 72 individuals were kept in one 90 litre glass tank (70 x 40 x 35 cm L x W x H) which contained approximately 100 to 150 individuals total. The tank contained artificial plants, plastic tunnels and a slow bubbling air stone. Water temperature was maintained at 25 ± 1°C (mean ± SD). Lighting was maintained on a 12:12 hr light:dark cycle. Fish were fed on a diet of live and fresh food (frozen blood worms, cyclops, mysis, brineshrimp and live banana worms) and ZM Granular pellets (© Copyright 2021 ZM Fish Food and Equipment).

### 3 Experimental Design

The experimental arena consisted of a five-arm maze constructed out of matt white corrugated PVC. Each arm (28 x 8 x 18 cm, L x W x H) was attached with white electrical tape to a 70 cm diameter circular base made of white acrylic. The experimental arena was suspended in a 83 x 63 cm (D x H) cylindrical tub with transparent 8 lb fishing wire attached to the base at five points to reduce vibrations (Figure 1a). A Sony AX53 digital 4K video camera recorder was suspended over the arena 161 cm above the base of the experimental arena (Figure 1b). The experimental arena was shielded from direct lighting and potential disturbance by white cotton sheets suspended around the arena and tub, while a black cotton sheet between the overhead fluorescent lighting and the camera reduced glare and reflections. The water level in the experimental arena was kept constant at 7.5 cm. Two Hepo HP-608 300W Aquarium heaters set to 25°C, one in the five-arm maze and one in the wider tub, and one Aquarium Systems Duetto 50 4W filter, were kept on between trials. The experimental arena was refilled every day before trials were started, and 50% of the water in the tub was replaced weekly. Water was allowed to fill gaps in the corrugated plastic walls of the experimental arena to prevent air pockets insulating sound passing through.

**Figure 1.**
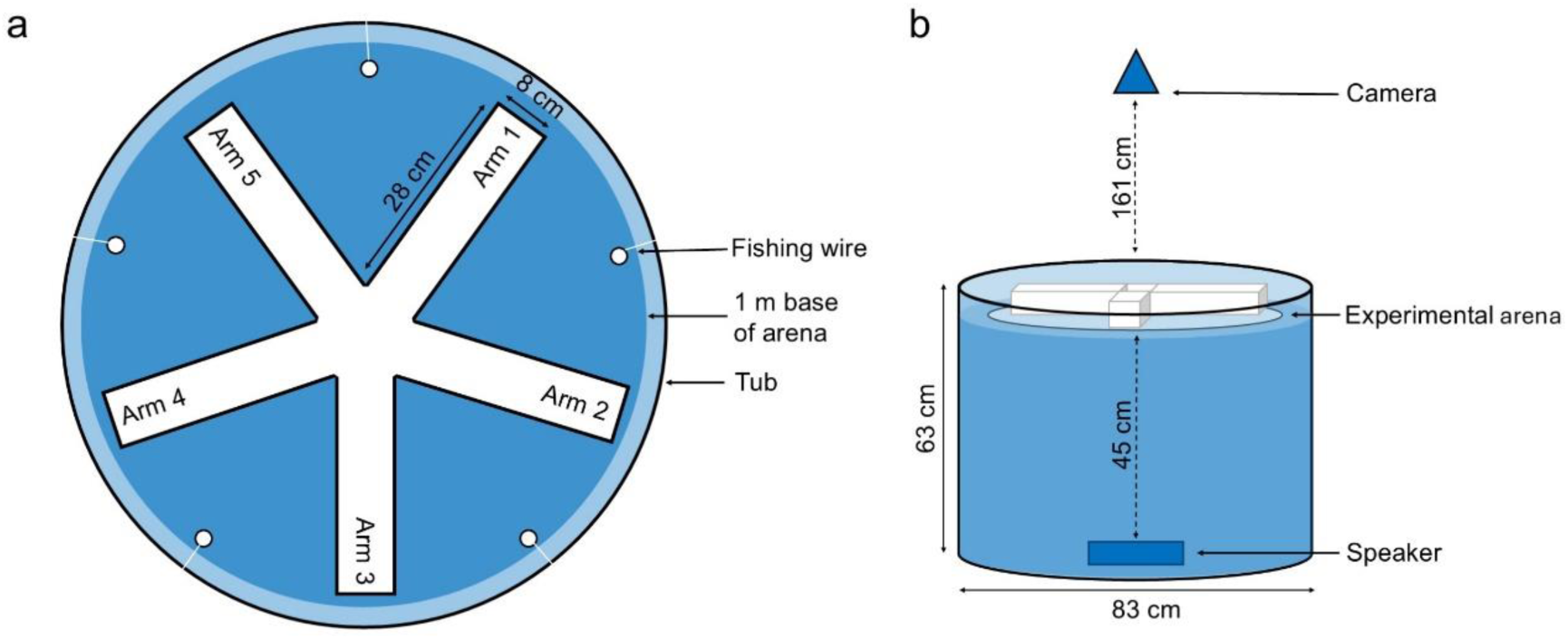
The experimental set up. (a) Overhead view of the experimental arena suspended in the tub by fishing wire. (b) Side view of experimental arena in tub with camera and loudspeaker placement. Not to scale.

### 4 Noise Treatments

An ambient playback track of sound recorded from the guppies’ housing tank was used as the control treatment, while the noise treatment consisted of white noise overlaid onto this ambient playback track (similar to Ginnaw *et al*., 2020). Only one track was used for each treatment (one for added white noise and one for the ambient control), as computer-generated standardised white noise exhibits minimal variation between tracks. White noise is characteristically consistent, and any minor variation is expected to be negligible, making it unlikely to introduce substantial variability between trials. The ambient playback track was recorded with a HiTech HTI-96-MIN omnidirectional hydrophone with inbuilt preamplifier and a Zoom H1n digital recorder (manufacturer-calibrated sensitivity: −164.3 dB re 1 V/mPa; frequency range 20e30 000 Hz, High Tech Inc., Gulfport, MS, U.S.A.) connected to a Zoom H6 digital recorder (PCMM10, 48 kHz sampling rate, Sony Corporation, Tokyo, Japan) in the housing tank at half water depth. A 5-minute recording was looped to generate a 30-minute track of ambient sound for the control treatment. The continuous white noise treatment was generated in Audacity (version 2.4.2) and overlaid onto the ambient sound track to simulate the effect of added anthropogenic noise on to their normal ambient soundscape.

Sounds recordings were made in both the housing tank and the experimental arena. Sound pressure recordings were made with a HiTech HTI-96-MIN omnidirectional hydrophone with inbuilt preamplifier connected to a Zoom H6 digital recorder. Reported sound levels are averages taken from recordings in each area of the experimental arena; the level was higher in the middle of the experimental arena because the small size of the arena caused the noise to concentrate more in the middle during playback (see Supplementary Table S1 and Figure S1 for more detailed analysis). The recorded RMS sound-pressure level in the guppies’ housing tank was 107 ± 2 dB re 1 μPa (mean ± SD), which the ambient control mimicked; the white noise treatment was distinctly louder at 135 ± 3 dB re 1 μPa. To measure the particle motion in the experimental arena, accelerometer recordings were taken using a M20-40 Geospectrum Technologies Inc. accelerometer and Zoom H6 digital recorder (see Supplementary Figure S2 for more detailed analysis). All acoustic recordings were analysed in MATLAB 2013a using paPAM (Nedelec *et al*., 2016). A bandpass filter was applied between 100 and 2000 Hz to account for the most common hearing sensitivities in fish (Popper and Fay, 2011). All heaters and filters were switched off during recordings, and water depth and temperature were kept constant, matching experimental arena conditions used during the trials.

Both playback tracks were 30 minutes in duration to provide continuous sound during the trial period. They were played from a SanDisk Clip Jam MP3 player through a DNH Aqua-30 underwater loudspeaker (frequency response 100–10,000 Hz, ADD DNH details) connected to an amplifier (Kemo Electronic GmbH; 18 W; frequency response: 40–20 000 Hz) and a Maplin 12 V 12 Ah battery. The loudspeaker was suspended in a plastic box using elastic to minimise vibrations and this was submerged in the bottom of the 63 x 83 cm black tub, 45 cm below the experimental arena (Figure 1b).

### 5 Experimental protocol

At the start of each week, groups of four female guppies (29.5 ± 2.4 mm, mean ± SD standard body length, *n* = 72 individuals) were formed from individuals haphazardly caught from the housing tank 24 hours before the start of the experimental trials. Individuals were randomly assigned to one of ten groups by shuffling the group numbers (from 1 to 10) randomly. Each fish was caught one at a time using a hand net and assigned to the corresponding group in the randomised order. The group numbers were then reshuffled, and the process repeated until four fish were in each group. Groups were formed in this way to minimise potential variation between groups, for example if bolder fish are caught first (King *et al*., 2013; MacGregor *et al*., 2020). Each group was held in a fry net (12.5 x 16 x 13.5 cm, L x W x H), with two nets per 45 litre tank (70 x 20 x 35 cm, L x W x H), in the same room as their original housing tank. Fish remained in these groups throughout the testing period of four days. Fish were fed ZM Granular pellets (© Copyright 2021 ZM Fish Food and Equipment) in the fry nets after being put into groups, and at the end of each day of trials. To control for differences in social and reproductive behaviour between males and females, only single-sex female groups were tested (Croft *et al*., 2003; Lucon-Xiccato *et al*., 2016).

All trials took place between 0900 and 1500 from the 20^th^ September to 5^th^ November 2021. Groups were tested once a day over four consecutive days in a repeated-measured design. The treatment that a group received alternated from one day to the next so all groups were tested twice in each treatment (noise or control), where a treatment (added noise) trial one day would be followed by a control trial the next day, and vice versa. Treatment order was decided by random shuffling of group numbers on the first day of testing: the first half of the random list were given the control treatment first and the second half given the noise treatment first. The testing order of groups within a day was also randomised. All randomisations were done using R (version 4.1.2; R Core Team, 2017).

At the start of a trial, all heaters and filters were turned off (pilot tests were conducted to ensure water temperature remained constant throughout the test day), the camera was switched on, and the appropriate playback track was started, to play through the underwater loudspeaker. A group of four fish were transferred to the experimental arena by a hand net and given a three-minute acclimation period inside a 12 (diameter) x 30 cm clear cylinder made of 5 mm rigid acrylic in the middle of the arena. At the end of the acclimation period, the cylinder was carefully lifted out of the arena with a clear rod attached to the cylinder with fishing wire to minimise disturbing the fish. After the 15-minute trial period, the camera filming was stopped, and the fish were caught and placed back into their fry net (unless it was the final day of testing, when they were released into a new housing tank for fish already used in the experiment). The heaters and filters were turned on for the period between the trials, and the experimental arena was set up for the next trial.

### 6 Video analysis and data processing

Video files were recorded in 4K (3840 x 2160 pixel) resolution with a frame rate of 25 fps. The automated tracking software idTracker (Pérez-Escudero *et al*., 2014) was run in MATLAB 2014a to obtain the trajectories of individual fish during the trials at each video frame. Identities were maintained within each trial but could not be confirmed across trials of the same group. Trajectories were then processed in R (version 4.1.2; R Core Team, 2017). In cases where there were missing coordinates for any individual in a frame, all data were removed from that frame. This is because with missing information for any individual, social parameters cannot be reliably calculated. Trajectories were smoothed using a Savitsky-Golay filter using the *Trajr* package in R (version 1.3.0; McLean and Skowron Volponi, 2018). Additionally, when the speed of an individual exceeded 25 pixels per frame, all data points for that frame were removed as this speed was assumed to be a result of erroneous tracking; cumulative density plots were used to determine the threshold at which high speeds were considered erroneous and removed. All trials were cropped to 13.5 minutes each, removing the first 1.5 minutes of the trial where the fish would often remain still after removal of the acclimation tube. Overall, tracked video footage for 72 trials across 18 groups was obtained.

### 7 Behavioural parameters

Using the processed trajectory data, the mean speed of each individual across each trial (pixels/frame) was calculated, and then the mean of these mean speeds across the four fish was calculated. Which arm each fish was in at every frame, or whether they were in the middle area joining the arms (Figure 1a), was determined using the coordinates of the corners of each arm, with these coordinates measured using ImageJ version 1.53 (Schneider *et al*., 2012). From this, four behavioural parameters were calculated per trial: mean cohesion (the standard deviation of the number of fish in each arm at each frame, averaged [mean] across all frames); the number of leadership attempts (moves into an empty arm); the number of following events (moves into an arm already occupied by another individual); and the total number of moves made into each arm (the sum of the leadership attempts and following events). During data processing, when distinguishing between whether movements into arms were leadership attempts or following events, four trials were found to have high rates of error. These four trials were removed from analyses relating to the number of leadership attempts or following events (sample size for these was 68 trials from 18 groups).

### 8 Statistical analysis

Mean speed, mean cohesion, and the total number of moves into arms (i.e. exploration) were analysed as response variables in linear mixed models (LMMs; constructed using the lmer function *lme4*, version 1.1.30; Bates *et al*., 2015). To analyse the proportion of moves into arms that were leadership attempts rather than following events, the leadership attempts and following events were transformed into a ratio using the cbind function in R (number of leadership attempts, number of following events) and this was used as the response variable in binomial generalised linear mixed models (GLMMs; constructed using the glmer function *lme4*, version 1.1.30; Bates *et al*., 2015). Based on the results from these models of the proportion of moves into arms that were leadership attempts, additional LMMs were constructed with the number of leadership attempts and the number of following events as separate response variables.

All models included group identity as a random effect and the time of day the trial began (trial start time) as a covariate. The inclusion of sound treatment (additional noise or ambient control) and day of testing (1 to 4) was varied between models to test for their independent and combined effects. For each response variable (i.e. mean speed, group cohesion, exploration, proportion of leadership attempts to following events, number of leadership attempts, number of following events), five models were constructed which included either the treatment*day interaction, treatment and day as main effects only, treatment only, day only, or neither of these terms as a null model. To determine which explanatory variables were important for explaining variation in the response, the Akaike Information Criterion values corrected for small samples sizes (AICc) were compared for each model using the ICtab function in R (bbmle version 1.0.24; Bolker *et al*., 2020). The model with a ΔAICc of zero is the most likely given the data. Models with a ΔAICc of greater than two units less than the null model were considered to have strong support, and therefore the fixed effects of the model were considered important in predicting the response variable (Burnham and Anderson, 2002; Symonds and Moussalli, 2011).

Individual identities were not tracked across trials, therefore repeatability was assessed at the group level only. To determine whether there was consistent variation among groups in the behavioural parameters over repeated tests, models with and without the random effect of group identity were compared for each response variable. Models included the fixed effects of start time and the interaction term of treatment*day. For continuous response variables, LMMs that included the random effect were compared to linear models (LMs) without it. For the binomial response variable (proportion of leadership attempts to following events), a binomial GLMM was compared to a binomial generalised linear model (GLM) without the random effect. ΔAICc values were used to evaluate the change in model likelihood when including the random effect for each response variable.

Consistent group-level repeatability was further tested using the *rpt* function in the *rptR* package (version 0.9.22; Stoffel et al., 2017) to calculate the intraclass correlation coefficient (ICC). Repeatability quantifies the proportion of total variance in a response variable that can be attributed to consistent differences between groups, relative to the total variance (including residual variance). Bootstrap resampling (1000 iterations) was applied to compute 95% confidence intervals for the repeatability estimates. We report the repeatability (R), 95% confidence intervals (CIs) and p-values from likelihood ratio tests (LRTs), with p < 0.05 indicating significant repeatability. The binomial GLMM for the proportion of leadership attempts to following events could not be used to calculate ICC or LRT values.

All analyses were conducted using R (version 4.1.2; R Core Team, 2017) in RStudio (R Core Team, 2017; RStudio Team, 2020). Multicollinearity was tested in all models by calculating the variation inflation factors (VIFs), with no strong evidence of multicollinearity found (VIF <3 in all cases). The assumption of normality in the residuals was tested in all LMMs using QQ plots, and residuals were plotted against fitted values to ensure homogeneity of variance. Maximum likelihood (ML) was used to fit all LMMs, rather than the *lme4* default restricted maximum likelihood (REML), because models within the comparisons contained different fixed effects (Faraway, 2005; Harrison *et al*., 2018). Under- or over-dispersion in all binomial GLMMs was tested using the residual diagnostics for mixed regression models (DHARMa; Hartig, 2019).

## Results

The model with testing day as the only explanatory variable was within 0.3 AICc units of the most likely model (with both sound treatment and day as main effects) and thus has similar predictive power. Including treatment did not substantially make the model more likely, suggesting that treatment did not affect mean speed (Table 1.1, Figure 2a). All models with day as a term were more likely (by more than 2 AICc units) than the null model, with the mean speed of individuals decreasing over the days of data collection (Figure 2a).

**Figure 2.**
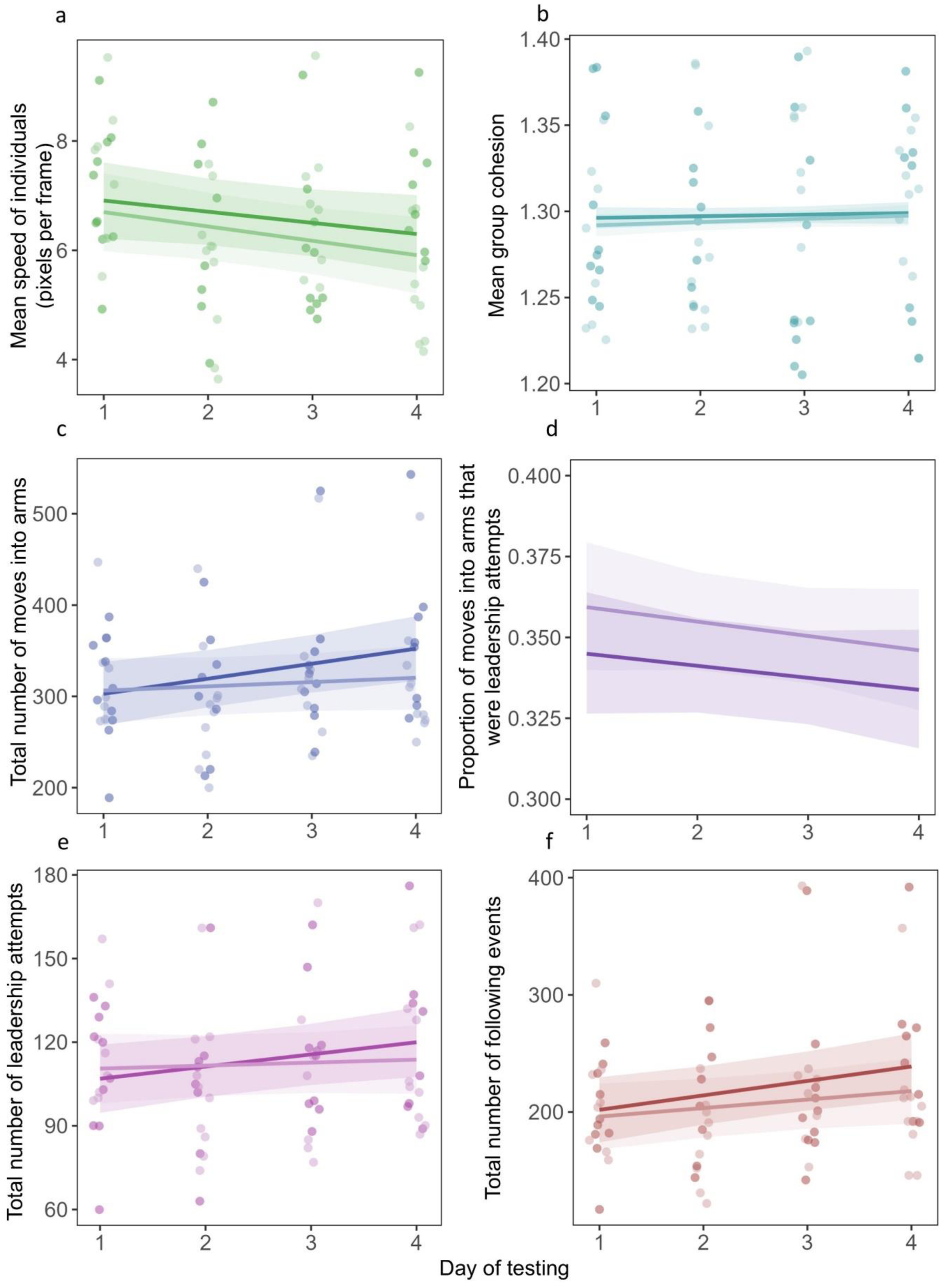
The relationship between the day of testing in both noise (lighter colour) and control (darker colour) treatments and the: (a) mean speed of individuals; (b) mean group cohesion (standard deviation of the number of fish in the same arm at any time); (c) total number of movements into arms during a trial; (d) proportion of moves into arms that were leadership events; (e) number of leadership attempts; and (f) number of following events. Fitted lines are calculated from fixed-effect estimates from the model with all main effects and the treatment*day interaction term. Points are individual data points (a, b and c: 18 groups, 72 trials; d, e and f: 18 groups, 68 trials) and are jittered (0.1 units) to reduce overplotting. The plot was generated using the *sjplot* package in r (Lüdecke *et al*., 2024).

**Table 1.**
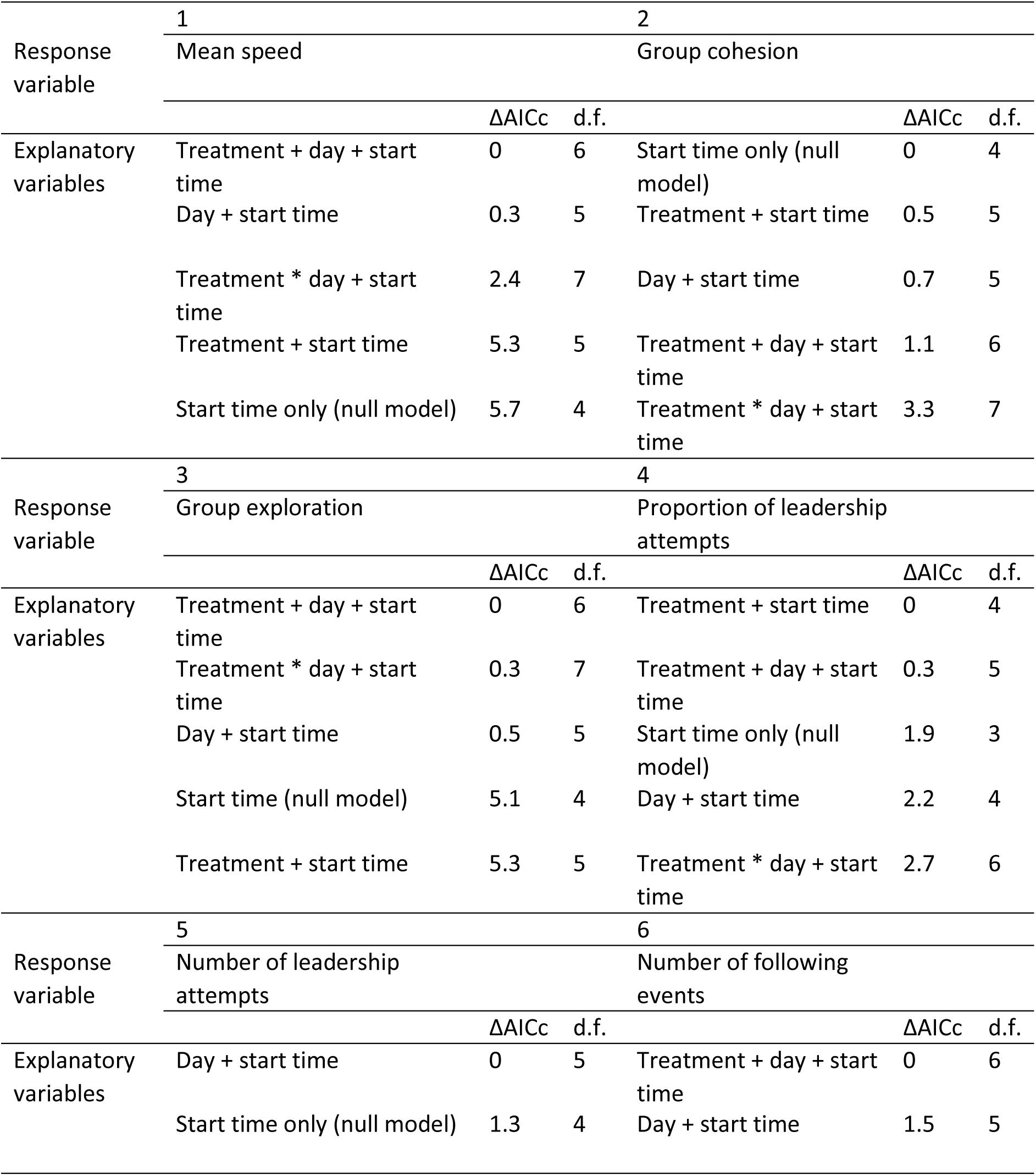
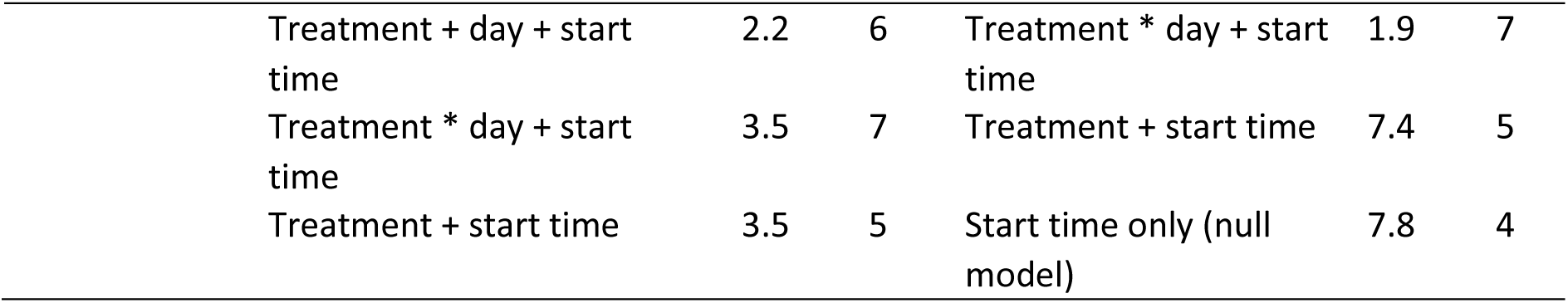
The ΔAICc for models explaining variation in the: (1) mean speed of individuals, (2) mean group cohesion, (3) group exploration (the total number of moves individuals made into arms), (4) proportion of moves into arms that were leadership attempts, (5) number of leadership attempts, and (6) number of following events. The null model includes start time as the only fixed effect. All models include group identity as the random effect. Treatment is noise or control, day is the day of testing (1 to 4).

When testing the variables that predicted the cohesion of fish during the trials, the most likely model was the null model (Table 1.2). All other models had larger AICc values than the null model so were not supported by the data (Figure 2b), suggesting no effects of treatment or testing day.

The analysis of exploration behaviour (total number of moves made into arms) suggested that including testing day as a main effect made the models more likely, as only models including day were substantially more likely than the null model, and the models that included treatment did not further improve the likelihood by more than 0.5 AICc units (Table 1.3). The model containing only treatment had a higher AICc value than the null model, also suggesting that treatment had no effect on the exploratory behaviour of groups. Groups were more exploratory as the days of testing progressed (Figure 2c).

When analysing the proportion of moves into arms that were leadership attempts, the most likely model had treatment as the explanatory variable (Table 1.4, Figure 2d). This model was 1.9 AICc units less than the null model, indicating moderate-to-strong evidence (Burnham and Anderson, 2002) that treatment affected the proportion of leadership attempts. There was a higher proportion of leadership attempts in the noise treatment compared to the control (Figure 2d). Including day as an additional predictor did not improve the model likelihood (AICc = 0.3 difference).

Given that in the previous analysis the difference in AICc values (1.9) was close to the threshold of 2 units that would indicate strong evidence that the proportion of leadership attempts was higher in the noise treatment, we explored this further by separately analysing the number of leadership attempts and following events. When the number of leadership attempts was the response variable, the most likely model was that including only day as an additional fixed effect. However, this was only 1.3 AICc units lower than the null model, indicating weak evidence that day affected the number of leadership attempts (Table 1.5, Figure 2d). All other models had AICc values higher than the null model and were therefore not supported.

For the number of following events as the response variable, the most likely model included both treatment and day as main effects (Table 1.6). The model without treatment, including the day and start time only, was 1.5 units from the most likely model, suggesting only moderate, rather than strong, support for an effect of treatment on the number of following events. Inclusion of testing day in the models improved their likelihood by >2 AICc units, suggesting strong evidence for day in predicting the number of following events. The total number of following events increased over the days of testing and was higher in the control treatment compared to the noise treatment (Figure 2e).

### Repeatability

For each response variable, models that included the random effect were more strongly supported by the data, with each having an AICc of zero and being more than 2 AICc units less than the model without the random effect (Table 2). This provides strong evidence that including group identity substantially improved the likelihood of the models and suggests there were consistent differences between groups (Figure 3). The R values further support this, with a significant proportion of the variation in behaviours (i.e. for mean speed, group cohesion, total moves into arms, leadership attempts and following events) being attributed to differences between groups (e.g. a repeatability of 0.676 indicates that 68% of the variation in speed was due to group-level differences). Similarly, exploration and the number of following events had high repeatability values of 0.744 and 0.519, respectively. Highly significant p-values (all <0.001) from the likelihood ratio tests reinforce the conclusion that repeatability is significantly greater than zero.

**Figure 3.**
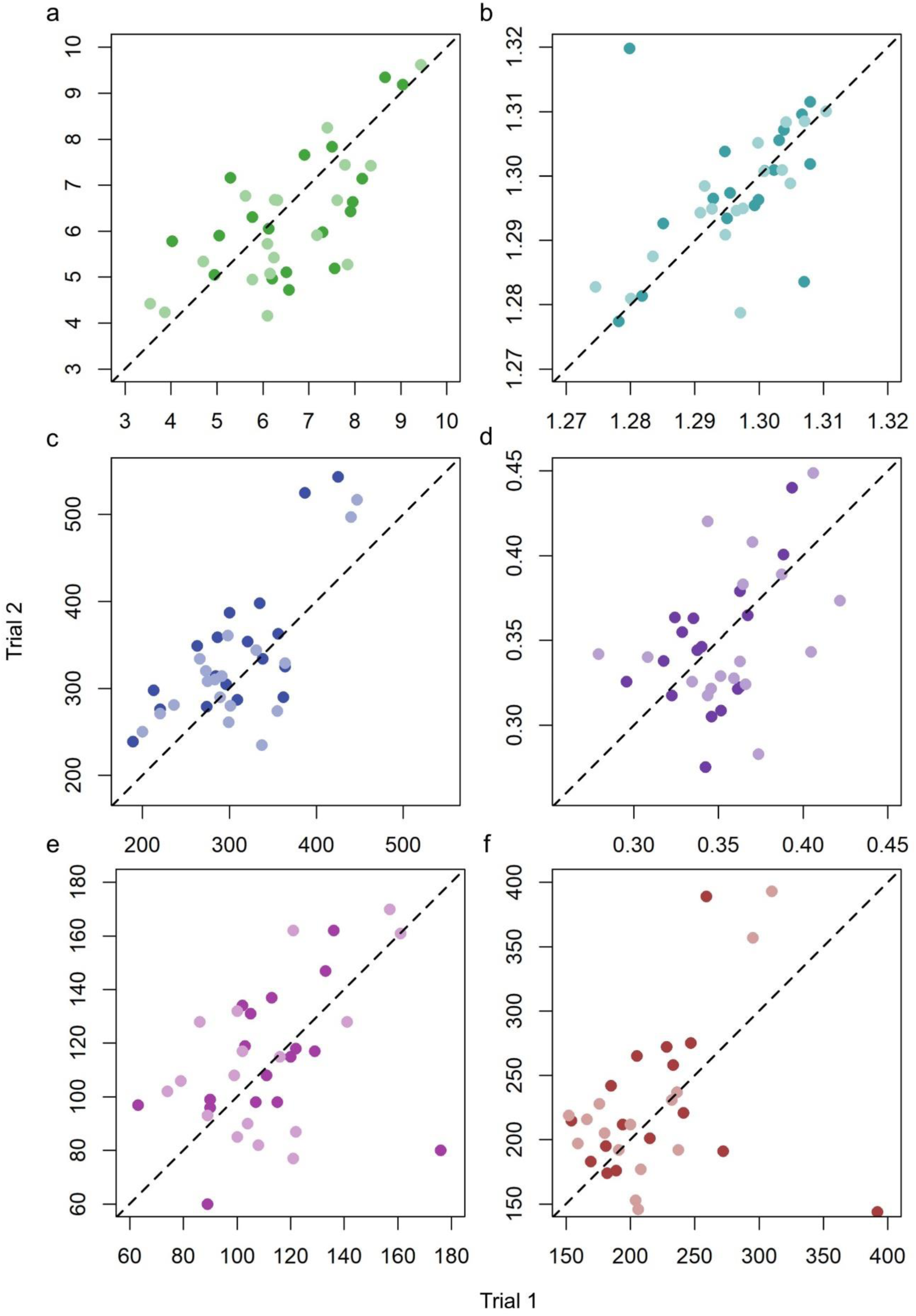
Relationships between response variables in the first and second control trials (darker points) and first and second treatment trials (lighter points) for each response variable: (a) mean speed of individuals (pixels per frame); (b) mean group cohesion (standard deviation of the number of fish in the same arm at any time); (c) total number of movements into arms during a trial; (d) proportion of moves into arms that were leadership events; (e) number of leadership attempts; and (f) number of following events. The dashed line is x = y.

**Table 2.** ΔAICc values, repeatability (R) and confidence intervals (CIs) from the intraclass correlation coefficient (ICC), and p-values from likelihood ratio tests (LRTs) for models with and without group identity as the random effect. All models include the fixed effects of start time and the interaction between treatment (noise or control) and day of testing (1 to 4). The binomial GLMM for the proportion of leadership attempts could not be used to calculate ICC or LRT values.

| Model | Random effect | ΔAICc | d.f. | R | 95% CI | p-value |
| --- | --- | --- | --- | --- | --- | --- |
| Mean speed ~<br>treatment * day + start time | Group ID | 0 | 7 | 0.676 | 0.431, 0.833 | <0.001 |
|  | NA | 35.8 | 6 |  |  |  |
| Group cohesion ~<br>treatment * day + start time | Group ID | 0 | 7 | 0.356 | 0.081, 0.604 | <0.001 |
|  | NA | 8 | 6 |  |  |  |
| Total moves into arms ~<br>treatment * day + start time | Group ID | 0 | 7 | 0.744 | 0.542, 0.867 | <0.001 |
|  | NA | 49.8 | 6 |  |  |  |
| Prop. leadership attempts ~<br>treatment * day + start time | Group ID | 0 | 6 | NA | NA | NA |
|  | NA | 26.6 | 5 |  |  |  |
| Total leadership attempts ~<br>treatment * day + start time | Group ID | 0 | 6 | 0.706 | 0.462, 0.846 | <0.001 |
|  | NA | 44.2 | 7 |  |  |  |
| Total following events ~<br>treatment * day + start time | Group ID | 0 | 6 | 0.776 | 0.58, 0.885 | <0.001 |
|  | NA | 69.6 | 7 |  |  |  |

## Discussion

The behaviour of animal groups under anthropogenic disturbances, such as noise pollution, is a growing field of study. Previous research has explored how shoaling behaviour is affected by various environmental factors, including temperature (Bartolini *et al*., 2015), turbidity (Chamberlain and Ioannou, 2019; MacGregor and Ioannou, 2023), dissolved oxygen (Domenici *et al*., 2002), dissolved carbon dioxide (Duteil *et al*., 2016), darkness (Miyazaki *et al*., 2000) and different combinations of multiple stressors (Ginnaw *et al*., 2020; Allibhai *et al*., 2023; Zanghi *et al*., 2023). Our results suggest that guppies may be more resilient to added acoustic noise than expected from previous studies (Currie *et al*., 2020, 2021; Fisher *et al*., 2021) as neither swimming speed nor group cohesion were affected by added white noise. There was some effect of white noise on the number of following events, as guppies in the noise treatment exhibited fewer following events than those in the control group. The most pronounced source of variation, however, was differences between groups, which showed high degrees of repeatability.

The absence of change in group cohesion as a result of increased acoustic noise was unexpected, as previous studies demonstrate that anthropogenic noise can reduce cohesion and coordination in shoals of juvenile seabass (*Dicentrarchus labrax*) under laboratory conditions (Herbert-Read *et al*., 2017b), and bluefin tuna (*Thunnus thynnus*) in the field (Sarà *et al*., 2007). Conversely, Eurasian minnows (*Phoxinus phoxinus*) were found to be more cohesive when exposed to added continuous sound (Currie *et al*., 2020; 2021). These studies used recordings of anthropogenic noise sources rather than white noise, which may yield different responses as a result of stress (Voellmy *et al*., 2014a). However, when white noise has been used in laboratory experiments, we observe similar results as found in other species, for example in three-spine sticklebacks (*Gasterosteus aculeatus*; Ginnaw *et al*., 2020), which showed that group cohesion was not impacted by the white noise treatment. Furthermore, in our study, group cohesion did not change over the days of testing. In studies of collective movement, experiments often find that over repeated trials, group cohesion reduces due to acclimation to experimental arenas (Miller and Gerlai, 2012; MacGregor and Ioannou, 2021). This is unlikely to be due to differences in the fish species used, as Allibhai *et al*. (2023) also used guppies and demonstrated this acclimation effect on group cohesion. Instead, the arena design may account for these differences, as our 5-arm radial maze was unlike the open arenas used in previous work (Allibhai *et al*., 2023; MacGregor and Ioannou, 2021; Miller and Gerlai, 2012).

There was some evidence that the proportion of all moves that were leadership events was higher in the noise treatment than in the control, with no change over the days of testing. Further examination revealed that this was because there were fewer following events in the noise treatment compared to the control, with no change in the number of leadership attempts. Added acoustic noise may be distracting individuals within a group from detecting social cues and following others (Purser and Radford, 2011; Voellmy *et al*., 2014a), although an effect of stress due to the added noise would be predicted to increase the tendency of fish to follow others (Herbert-Read et al., 2019). The movements of other fish in the group are critical for collective movement and decision-making (Ioannou *et al*., 2011; Lemasson *et al*., 2018), and are a form of social information; our results suggest that noise could be restricting the ability of group members to use social information, and causing them to be unresponsive to leadership cues. This could have consequences for the fitness benefits provided by group-living, where the use of social information is key to making better decisions. However, this reduction in followership did not have a detectable impact on group cohesion as this was not affected by added acoustic noise, thus the fitness effects of reduced followership deserves further study.

The most pronounced source of variation in the swimming speed and collective behaviour variables was the consistent differences among the groups tested over multiple days. These consistent, repeatable differences among groups across all behavioural parameters is considered group-level personality variation, as found in previous research (Planas-Sitjà *et al*., 2015; Salazar *et al*., 2015; Jolles *et al*., 2018; MacGregor and Ioannou, 2021). This group-level personality variation may arise from stable individual differences within the group or emerge from interactions between group members (Jolles *et al*., 2017). High variability in group personality could influence how susceptible or resilient a group is to external disturbances, suggesting that personality traits might buffer groups from potential stressors (Harding *et al*., 2019). However, with only two replicates per group per treatment and only two levels (control and noise treatment), our study was not designed to explore this, but this would likely be a fruitful avenue for future research.

Our results reflect recent findings on a major predator of guppies, pike cichlids. In the study of Brown and Ioannou (2024), feeding motivation, and hence the risk they likely pose to their prey, of the pike cichlid *Saxatilia proteus* was found to be unchanged by environmental parameters (temperature and light), but showed strong among individual variation that was consistent over time. The importance of personality variation in the risk that pike cichlids pose has also been demonstrated in the natural habitat of guppies (Szopa-Comley et al., 2020). Together with our current study, the next step is to explore the interaction between the personality types of individual pike cichlids and personality types of small guppy shoals, for example testing whether the success of the predators or prey is dependent more on the personality type of one than the other, or is dependent on the interaction between the personality types of both predator and prey.

Anthropogenic noise is increasingly recognised as a significant disturbance in aquatic ecosystems, where masking, distraction and stress generate unimodal or cross-modal effects on animal behaviour (Slabbekoorn et al., 2010; Duarte et al., 2021). In group-living species, these disturbances may destabilise group structure and reduce cohesion, both of which are vital for maintaining the benefits provided by group-living, such as improved predator avoidance (Krause & Ruxton, 2002; Ioannou, 2021). Our study examined how decision-making in guppy shoals responds to added white noise and found that, overall, guppies were robust to this environmental stressor in the behaviours examined. However, a subtle reduction in following behaviour in the noise treatment suggests that white noise may disrupt social decision-making processes. This finding underscores the importance of examining specific social behaviours, like decision-making. Understanding these nuances will be crucial as we further explore how environmental stressors influence group-living species and their complex social structures.

## Acknowledgements

This work was supported by Natural Environment Research Council grant NE/P012639/1 awarded to C.C.I. We would like to thank Emma Weschke, Isla Keesje Davidson and Josh W. Pysanczyn for providing advice in recording underwater audio; Josh W. Pysanczyn for guidance on setting up underwater playback equipment and generating audio files; Sophie L. Nedelec for training in analysing audio data; and Costanza Zanghí for advice on the manuscript.

## Author contributions

M.A.C. developed the experimental design, conducted data collection and data analysis, and wrote the first draft of the manuscript; M.A.C. and E.W. conducted noise recordings and analysed the audio data. A.N.R and S.D.S supervised M.A.C and E.W. in designing noise treatments and analysing audio data. C.C.I. conceived the project and supervised M.A.C. and E.W. All authors reviewed the manuscript.

## Data accessibility

We have included our data and code in this submission, which will be uploaded to an online repository upon acceptance.

## References

Allibhai, I., Zanghi, C., How, M. J., & Ioannou, C. C. (2023). Increased water temperature and turbidity act independently to alter social behavior in guppies (Poecilia reticulata). Ecology and Evolution, 13(3), e9958. 10.1002/ece3.9958

Bartolini, T., Butail, S., & Porfiri, M. (2015). Temperature influences sociality and activity of freshwater fish. Environmental Biology of Fishes, 98(3), 825–832. 10.1007/s10641-014-0318-8

Bates, D., Mächler, M., Bolker, B., & Walker, S. (2015). Fitting linear mixed-effects models using lme4. Journal of Statistical Software, 67, 1–48. 10.18637/jss.v067.i01

Bolker, B., Team, R. D. C., & Giné-Vázquez, I. (2020). bbmle: Tools for General Maximum Likelihood Estimation (Version 1.0.25) [Computer software]. https://CRAN.R-project.org/package=bbmle

Brown, L. J., & Ioannou, C. C. (2024). Predator Personality Variation, Not the Multiple Stressors of Temperature and Light, Determines Feeding Motivation in an Ambush Piscivore Saxatilia proteus. Ecology and Evolution, 14(11), e70540. 10.1002/ece3.70540

Bumann, D., & Krause, J. (1993). Front individuals lead in shoals of three-spined sticklebacks (Gasterosteus aculeatus) and juvenile roach (Rutilus rutilus). Behaviour, 125, 189–198. 10.1163/156853993X00236

Burnham, K. P., & Anderson, D. R. (2002). Model Selection and Inference: A Practical Information-Theoretic Approach. 2nd Edition. Springer.

Chamberlain, A. C., & Ioannou, C. C. (2019). Turbidity increases risk perception but constrains collective behaviour during foraging by fish shoals. Animal Behaviour, 156, 129–138. 10.1016/j.anbehav.2019.08.012

Conradt, L., & Roper, T. J. (2005). Consensus decision making in animals. Trends in Ecology & Evolution, 20(8), 449–456. 10.1016/j.tree.2005.05.008

Croft, D. P., Arrowsmith, B. J., Bielby, J., Skinner, K., White, E., Couzin, I. D., Magurran, A. E., Ramnarine, I., & Krause, J. (2003). Mechanisms underlying shoal composition in the Trinidadian guppy, Poecilia reticulata. Oikos, 100(3), 429–438.

Currie, H. A. L., White, P. R., Leighton, T. G., & Kemp, P. S. (2020). Group behavior and tolerance of Eurasian minnow (Phoxinus phoxinus) in response to tones of differing pulse repetition rate. The Journal of the Acoustical Society of America, 147(3), 1709– 1718. 10.1121/10.0000910

Currie, H. A. L., White, P. R., Leighton, T. G., & Kemp, P. S. (2021). Collective behaviour of the European minnow (*Phoxinus phoxinus*) is influenced by signals of differing acoustic complexity. Behavioural Processes, 189, 104416. 10.1016/j.beproc.2021.104416

Domenici, P., Silvana Ferrari, R., Steffensen, J. F., & Batty, R. S. (2002). The effect of progressive hypoxia on school structure and dynamics in Atlantic herring Clupea harengus. Proceedings of the Royal Society of London. Series B: Biological Sciences, 269(1505), 2103–2111. 10.1098/rspb.2002.2107

Duarte, C. M., Chapuis, L., Collin, S. P., Costa, D. P., Devassy, R. P., Eguiluz, V. M., Erbe, C., Gordon, T. A. C., Halpern, B. S., Harding, H. R., Havlik, M. N., Meekan, M., Merchant, N. D., Miksis-Olds, J. L., Parsons, M., Predragovic, M., Radford, A. N., Radford, C. A., Simpson, S. D.,… Juanes, F. (2021). The soundscape of the Anthropocene ocean. Science (New York, N.Y.), 371(6529), eaba4658. 10.1126/science.aba4658

Duteil, M., Pope, E. C., Pérez-Escudero, A., de Polavieja, G. G., Fürtbauer, I., Brown, M. R., & King, A. J. (2016). European sea bass show behavioural resilience to near-future ocean acidification. Royal Society Open Science, 3(11), 160656. 10.1098/rsos.160656

Faraway, J. J. (2005). Extending the Linear Model with R: Generalized Linear, Mixed Effects and Nonparametric Regression Models (1st edition). Chapman and Hall/CRC.

Fisher, D. N., Kilgour, R. J., Siracusa, E. R., Foote, J. R., Hobson, E. A., Montiglio, P.-O., Saltz, J. B., Wey, T. W., & Wice, E. W. (2021). Anticipated effects of abiotic environmental change on intraspecific social interactions. Biological Reviews, 96(6), 2661–2693. 10.1111/brv.12772

Foote, A. D., Osborne, R. W., & Hoelzel, A. R. (2004). Whale-call response to masking boat noise. Nature, 428, 910–910. 10.1038/428910a

Ginnaw, G. M., Davidson, I. K., Harding, H. R., Simpson, S. D., Roberts, N. W., Radford, A. N., & Ioannou, C. C. (2020). Effects of multiple stressors on fish shoal collective motion are independent and vary with shoaling metric. Animal Behaviour, 168, 7–17. 10.1016/j.anbehav.2020.07.024

Harding, H. R., Gordon, T. A. C., Eastcott, E., Simpson, S. D., & Radford, A. N. (2019). Causes and consequences of intraspecific variation in animal responses to anthropogenic noise. Behavioral Ecology, 30(6), 1501–1511. 10.1093/beheco/arz114

Harrison, X. A., Donaldson, L., Correa-Cano, M. E., Evans, J., Fisher, D. N., Goodwin, C. E. D., Robinson, B. S., Hodgson, D. J., & Inger, R. (2018). A brief introduction to mixed effects modelling and multi-model inference in ecology. PeerJ, 6, e4794. 10.7717/peerj.4794

Hartig, F. (2019). DHARMa: Residual diagnostics for hierarchical (multi-level / mixed) regression models (R package version 0.2.6.).

Herbert-Read, J. E., Kremer, L., Bruintjes, R., Radford, A. N., & Ioannou, C. C. (2017). Anthropogenic noise pollution from pile-driving disrupts the structure and dynamics of fish shoals. Proceedings of the Royal Society B: Biological Sciences, 284(1863). 10.1098/rspb.2017.1627

Herbert-Read, J. E., Wade, A. S. I., Ramnarine, I. W., & Ioannou, C. C. (2019). Collective decision-making appears more egalitarian in populations where group fission costs are higher. Biology Letters, 15(12), 20190556. 10.1098/rsbl.2019.0556

Ioannou, C. (2021). Grouping and Predation. In T. K. Shackelford & V. A. Weekes-Shackelford (Eds.), Encyclopedia of Evolutionary Psychological Science (pp. 3574–3580). Springer International Publishing. 10.1007/978-3-319-19650-3_2699

Ioannou, C. C., Couzin, I. D., James, R., Croft, D. P., & Krause, J. (2011). Social organisation and information transfer in schooling fish. In Fish cognition and behaviour (2nd ed., pp. 217–239).

Ioannou, C. C., & Laskowski, K. L. (2023a). A multi-scale review of the dynamics of collective behaviour: From rapid responses to ontogeny and evolution. Philosophical Transactions of the Royal Society B: Biological Sciences, 378(1874), 20220059. 10.1098/rstb.2022.0059

Ioannou, C. C., & Laskowski, K. L. (2023b). Conformity and differentiation are two sides of the same coin. Trends in Ecology & Evolution, 38(6), 545–553. 10.1016/j.tree.2023.01.014

Ioannou, C. C., Ramnarine, I. W., & Torney, C. J. (2017). High-predation habitats affect the social dynamics of collective exploration in a shoaling fish. Science Advances, 3(5), e1602682. 10.1126/sciadv.1602682

Jolles, J. W., Boogert, N. J., Sridhar, V. H., Couzin, I. D., & Manica, A. (2017). Consistent Individual Differences Drive Collective Behavior and Group Functioning of Schooling Fish. Current Biology, 27(18), 2862–2868.e7. 10.1016/j.cub.2017.08.004

Jolles, J. W., Laskowski, K. L., Boogert, N. J., & Manica, A. (2018). Repeatable group differences in the collective behaviour of stickleback shoals across ecological contexts. Proceedings of the Royal Society B: Biological Sciences, 285(1872), 20172629. 10.1098/rspb.2017.2629

King, A. J. (2010). Follow me! I’m a leader if you do; I’m a failed initiator if you don’t? Behavioural Processes, 84(3), 671–674. 10.1016/j.beproc.2010.03.006

King, A. J., Fürtbauer, I., Mamuneas, D., James, C., & Manica, A. (2013). Sex-differences and temporal consistency in stickleback fish boldness. PLOS ONE, 8(12), e81116. 10.1371/journal.pone.0081116

Krause, J., Hoare, D., Krause, S., Hemelrijk, C. K., & Rubenstein, D. I. (2000). Leadership in fish shoals. Fish and Fisheries, 1(1), 82–89. 10.1111/j.1467-2979.2000.tb00001.x

Krause, J., & Ruxton, G. D. (2002). Living in Groups. Oxford University Press. https://books.google.co.uk/books?hl=en&lr=&id=HAoUFfVFtMcC&oi=fnd&pg=PA1&ots=mn3JhH89e4&sig=AZVbJX19WXA59XEQ0-H8DKKaYVQ&redir_esc=y#v=onepage&q&f=false

Lemasson, B., Tanner, C., Woodley, C., Threadgill, T., Qarqish, S., & Smith, D. (2018). Motion cues tune social influence in shoaling fish. Scientific Reports, 8(1), Article 1. 10.1038/s41598-018-27807-1

Lucon-Xiccato, T., Dadda, M., & Bisazza, A. (2016). Sex differences in discrimination of shoal size in the guppy (Poecilia reticulata). Ethology, 122(6), 481–491. 10.1111/eth.12498

Lüdecke, D., Bartel, A., Schwemmer, C., Powell, C., Djalovski, A., & Titz, J. (2024). sjPlot: Data Visualization for Statistics in Social Science (Version 2.8.16) [Computer software]. https://cran.r-project.org/web/packages/sjPlot/index.html

MacGregor, H. E. A., Herbert-Read, J. E., & Ioannou, C. C. (2020). Information can explain the dynamics of group order in animal collective behaviour. Nature Communications, 11(1). 10.1038/s41467-020-16578-x

MacGregor, H. E. A., & Ioannou, C. C. (2021). Collective motion diminishes, but variation between groups emerges, through time in fish shoals. Royal Society Open Science, 8(10), 210655. 10.1098/rsos.210655

MacGregor, H. E. A., & Ioannou, C. C. (2023). Shoaling behaviour in response to turbidity in three-spined sticklebacks. Ecology and Evolution, 13(11), e10708. 10.1002/ece3.10708

McLean, D. J., & Skowron Volponi, M. A. (2018). trajr: An R package for characterisation of animal trajectories. Ethology, 124(6), 440–448. 10.1111/eth.12739

Miller, N., & Gerlai, R. (2012). From schooling to shoaling: Patterns of collective motion in Zebrafish (Danio rerio). PLOS ONE, 7(11), e48865. 10.1371/journal.pone.0048865

Miyazaki, T., Shiozawa, S., Kogane, T., Masuda, R., Maruyama, K., & Tsukamoto, K. (2000). Developmental changes of the light intensity threshold for school formation in the striped jack Pseudocaranx dentex. Marine Ecology Progress Series, 192, 267–275. 10.3354/meps192267

Morris-Drake, A., Kern, J. M., & Radford, A. N. (2016). Cross-modal impacts of anthropogenic noise on information use. Current Biology, 26(20), R911–R912. 10.1016/j.cub.2016.08.064

Nedelec, S. L., Campbell, J., Radford, A. N., Simpson, S. D., & Merchant, N. D. (2016). Particle motion: The missing link in underwater acoustic ecology. Methods in Ecology and Evolution, 7(7), 836–842. 10.1111/2041-210X.12544

Parks, S. E., Clark, C. W., & Tyack, P. L. (2007). Short- and long-term changes in right whale calling behavior: The potential effects of noise on acoustic communication. The Journal of the Acoustical Society of America, 122(6). 10.1121/1.2799904

Peng, C., Zhao, X., & Liu, G. (2015). Noise in the sea and its impacts on marine organisms. International Journal of Environmental Research and Public Health. 10.3390/ijerph121012304

Pérez-Escudero, A., Vicente-Page, J., Hinz, R. C., Arganda, S., & de Polavieja, G. G. (2014). idTracker: Tracking individuals in a group by automatic identification of unmarked animals. Nature Methods, 11(7), Article 7. 10.1038/nmeth.2994

Planas-Sitjà, I., Deneubourg, J.-L., Gibon, C., & Sempo, G. (2015). Group personality during collective decision-making: A multi-level approach. Proceedings of the Royal Society B: Biological Sciences, 282(1802), 20142515. 10.1098/rspb.2014.2515

Popper, A. N., & Fay, R. R. (2011). Rethinking sound detection by fishes. Hearing Research, 273(1–2), 25–36. 10.1016/j.heares.2009.12.023

Purser, J., & Radford, A. N. (2011). Acoustic noise induces attention shifts and reduces foraging performance in three-spined sticklebacks (Gasterosteus aculeatus). PLoS ONE. 10.1371/journal.pone.0017478

R Core Team. (2017). R: A language and environment for statistical computing. R Foundation for Statistical Computing, Vienna, Austria. URL https://www.R-project.org/. [Computer software].

RStudio Team. (2020). RStudio: Integrated Development for R. [Computer software]. RStudio, PBC, Boston, MA.

Salazar, M.-O. L., Planas-Sitjà, I., Deneubourg, J.-L., & Sempo, G. (2015). Collective resilience in a disturbed environment: Stability of the activity rhythm and group personality in Periplaneta americana. Behavioral Ecology and Sociobiology, 69(11), 1879–1896.

Sarà, G., Dean, J., D’ Amato, D., Buscaino, G., Oliveri, A., Genovese, S., Ferro, S., Buffa, G., Martire, M., & Mazzola, S. (2007). Effect of boat noise on the behaviour of bluefin tuna Thunnus thynnus in the Mediterranean Sea. Marine Ecology Progress Series, 331, 243–253. 10.3354/meps331243

Schneider, C. A., Rasband, W. S., & Eliceiri, K. W. (2012). NIH Image to ImageJ: 25 years of image analysis. Nature Methods, 9(7), 671–675. 10.1038/nmeth.2089

Slabbekoorn, H., Bouton, N., van Opzeeland, I., Coers, A., ten Cate, C., & Popper, A. N. (2010). A noisy spring: The impact of globally rising underwater sound levels on fish. Trends in Ecology & Evolution, 25(7), 419–427. 10.1016/J.TREE.2010.04.005

Smith, M. E., Kane, A. S., & Popper, A. N. (2004). Noise-induced stress response and hearing loss in goldfish (Carassius auratus). Journal of Experimental Biology, 207(3), 427–435. 10.1242/jeb.00755

Sumpter, D. j. t. (2006). The principles of collective animal behaviour. Philosophical Transactions of the Royal Society B: Biological Sciences, 361(1465), 5–22. 10.1098/rstb.2005.1733

Symonds, M. R. E., & Moussalli, A. (2011). A brief guide to model selection, multimodel inference and model averaging in behavioural ecology using Akaike’s information criterion. Behavioral Ecology and Sociobiology, 65(1), 13–21. 10.1007/s00265-010-1037-6

Szopa-Comley, A. W., Duffield, C., Ramnarine, I. W., & Ioannou, C. C. (2020). Predatory behaviour as a personality trait in a wild fish population. Animal Behaviour, 170, 51–64. 10.1016/j.anbehav.2020.10.002

Voellmy, I. K., Purser, J., Flynn, D., Kennedy, P., Simpson, S. D., & Radford, A. N. (2014). Acoustic noise reduces foraging success in two sympatric fish species via different mechanisms. Animal Behaviour, 89, 191–198. 10.1016/j.anbehav.2013.12.029

Wysocki, L. E., Dittami, J. P., & Ladich, F. (2006). Ship noise and cortisol secretion in European freshwater fishes. Biological Conservation, 128(4), 501–508. 10.1016/j.biocon.2005.10.020

Zanghi, C., Munro, M., & Ioannou, C. C. (2023). Temperature and turbidity interact synergistically to alter anti-predator behaviour in the Trinidadian guppy. Proceedings of the Royal Society B: Biological Sciences, 290(2002), 20230961. 10.1098/rspb.2023.0961

