## Supplementary table 1, figure 1 and figure 2 for "Group personality, rather than acoustic noise, causes variation in group decision-making in guppy shoals"

**Table S1.** The RMS noise level in dB re 1 μPa for each area of the experimental arena reported as mean ± standard deviation. Hydrophone recordings were taken from the experimental arena for both treatments using a HiTech HTI-96-MIN hydrophone and Zoom H1n digital recorder (sensitivity levels: 3 for ambient playback, 5 for white noise playback). 10-second recordings were taken at the end, middle, and start of each arm, and one recording taken in the middle of the experimental arena. S1 refers to the corresponding letters in Figure S1. Hydrophone recordings were analysed in MATLAB v2013a using paPAM (Nedelec *et al.,* 2016).

| S1 | Area of experimental arena | Ambient control treatment (RMS level in dB re 1 μPa) | Added white noise treatment (RMS level in dB re 1 μPa) |
|---|---|---|---|
| A | Middle of arena | 112 | 138 |
| B | Start of arms | 107 ± 0.9 | 136 ± 1.0 |
| C | Middle of arms | 106 ± 1.2 | 135 ± 1.0 |
| D | End of arms | 105 ± 0.6 | 132 ± 0.7 |

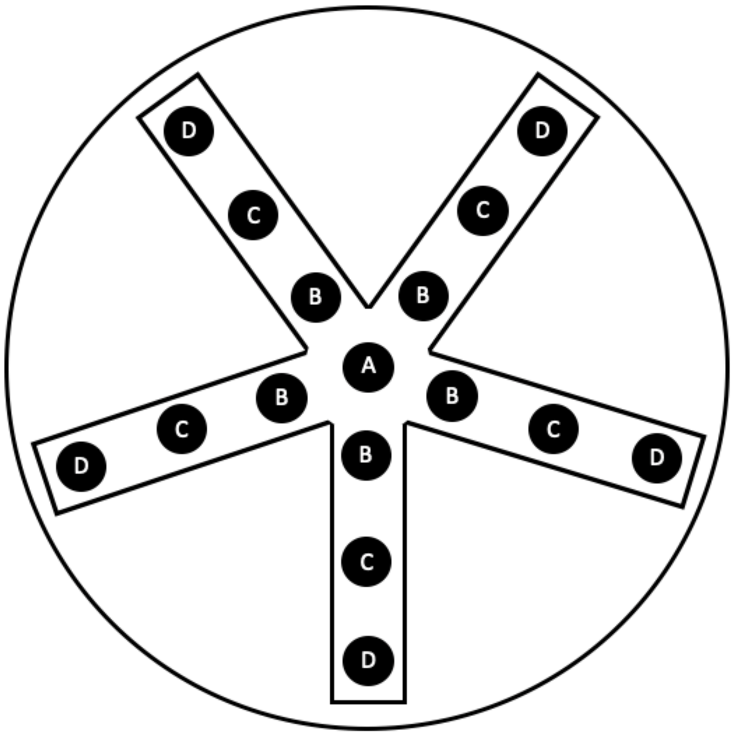


**Figure S1.** Areas of experimental arena where hydrophone recordings were taken, corresponding to data in Table S1.

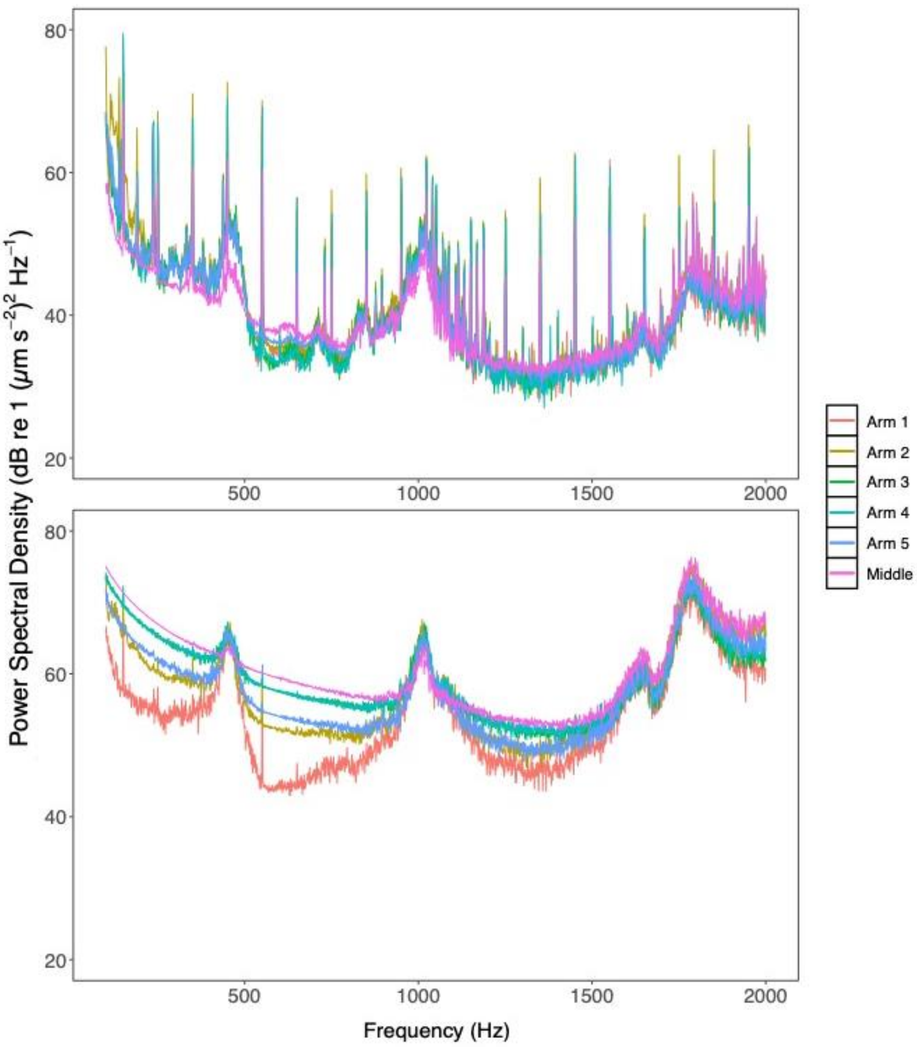


**Figure S2.** Power spectral density plot for particle motion in each area of the arena for both the ambient control treatment (top) and white noise treatment (bottom). Recordings were taken using a M20-40 Geospectrum Technologies Inc. accelerometer and Zoom H6 digital recorder (sensitivity level 3 for ambient, 5 for white noise playback). Recordings were made in middle of the experimental arena and once in each arm. Due to the shallow water depth and relatively large size of the accelerometer, it had to be orientated horizontally and thus could only record on the Z axis. 30-second recordings were taken and these were cropped to 10 seconds for analysis. The mean values for particle motion throughout the experimental arena were 30.58 – 81.56 dB re 1 μm $s^{-2}$ in the ambient control treatment and 49.66 – 97.99 dB re 1 μm s−2 in the added white noise treatment.
